# *Staphylococcus epidermidis* enables *Cutibacterium acnes* to form biofilms under aerobic conditions

**DOI:** 10.1101/2023.10.12.562081

**Authors:** Jeffrey B. Kaplan

**Affiliations:** Department of Biology, American University, Washington DC 20016, USA

**Keywords:** Acne vulgaris, Aerobic, Anaerobic, Biofilm, Dispersin B, DNase, Extracellular DNA, Implant infection, PNAG

## Abstract

*Staphylococcus epidermidis* and *Cutibacterium acnes* are among the most abundant members of the human skin microbiome. Both species are associated with skin health and disease. Although skin microbes typically grow in surface-associated biofilms, few studies on the interaction between *S. epidermidis* and *C. acnes* in biofilms have been reported. In the present study we measured the ability of *S. epidermidis* and *C. acnes*, either individually or jointly, to form biofilms in glass culture tubes. Since *S. epidermidis* is a facultative anaerobe and *C. acnes* is an aerotolerant anaerobe, tubes were incubated both aerobically and anaerobically to assess the role of atmosphere in biofilm forma4on. When cultured individually, we found that *S. epidermidis* formed biofilms under both aerobic and anaerobic conditions, whereas *C. acnes* formed biofilms only under aerobic conditions. When co-cultured, the presence of *C. acnes* had no effect on *S. epidermidis* biofilm formation under either aerobic or anaerobic conditions. However, the presence of *S. epidermidis* significantly enhanced the growth of *C. acnes* biofilms under anaerobic conditions and enabled *C. acnes* to form biofilms under aerobic condi2ons. This finding may be clinically relevant to the interaction between *S. epidermidis* and *C. acnes* on human skin.

## INTRODUCTION

The Gram-positive bacteria *Staphylococcus epidermidis* and *Cutibacterium acnes* are among the most abundant members of the human skin microbiome (Ahle *et al*., 2022; Fournière *et al*., 2020). Although these species are considered beneficial because they maintain homeostasis of the skin microbiota and control pathogens such as *Staphylococcus. aureus* via colonization resistance (Francuzik *et al*., 2018; Rozas *et al*., 2021; Severn & Horswill, 2023), both species can also act as opportunis)c pathogens. *S. epidermidis* is associated with bacteremia in preterm infants and atopic dermatitis (Severn & Horswill, 2023), whereas C. *acnes* can cause invasive infections of the skin, soft tissue, and cardiovascular system (Achermann *et al*., 2014). Both species also cohabitate in acne vulgaris lesions (McLaughlin *et al*., 2019) and cause infections of implanted medical devices (Achermann *et al*., 2014; Otto, 2009).

Biofilms are defined as densely packed layers of bacterial cells growing atiached to a tissue or surface (Costerton *et al*., 1999). Bacteria in a biofilm are encased in a sticky, self-synthesized, extracellular polymeric matrix that holds the cells together in a mass, attaches them to the underlying surface, and protects them from killing by an1microbial agents and host immunity. Biofilms play a role in many chronic infections which are often difficult to treat because of the protective nature biofilms. *In vivo, S. epidermidis* biofilms have been observed on skin, in sweat glands, and on the surfaces of implanted medical devices (Allen & Mueller, 2011; Büttner *et al*., 2015), and *C. acnes* biofilms have been observed in acne lesions and on and implanted medical devices (Bayston *et al*., 2006; Jahns *et al*., 2012). Both *S. epidermidis* and *C. acnes* have been shown to form biofilms *in vitro* (Achermann *et al*., 2014; Otto, 2009).

Since *S. epidermidis* and *C. acnes* are both abundant colonizers of human skin, these species likely evolved a close relationship based on competition and mutualism. For example, previous studies showed that *C. acnes* competes with *S. epidermidis* in hair follicles through the production of the antimicrobial peptide cutimycin (Claesen *et al*., 2020). *C. acnes* may also inhibit *S. epidermidis* biofilm formation and sensitizes *S. epidermidis* to antibiotic killing through the production of short-chain fatty acids (Nakamura *et al*., 2020). *S. epidermidis*, however, can counteract this competition through the production of antimicrobial peptides and the fermentation of glycerol into short-chain fatty acids which suppress *C. acnes* growth (Wang *et al*., 2014). In addition, *S. epidermidis* and *C. acnes* likely evolved the ability to coexist in microbial biofilms on human skin (Brandwein *et al*., 2016), although few studies investigating the interaction between *S. epidermidis* and *C. acnes* in dual-species or multi-species biofilms have been reported.

In the present study we assessed the ability of *S. epidermidis* and *C. acnes*, either individually or jointly, to form biofilms in glass culture tubes. Since *S. epidermidis* is a facultative anaerobe and *C. acnes* is an aerotolerant anaerobe, we incubated the tubes under both aerobic and anaerobic conditions to determine the role of atmosphere in biofilm formation. Our major finding was that *S. epidermidis* significantly enhanced the growth of *C. acnes* biofilms under anaerobic conditions and enabled *C. acnes* to form biofilms under aerobic conditions. This finding may be clinically relevant to the interaction between *S. epidermidis* and *C. acnes* on human skin.

## MATERIALS AND METHODS

### Bacterial strains and growth conditions

The bacterial strains used in this study are listed in Table 1. Bacteria were maintained and enumerated on Trypttc Soy agar. Biofilms were cultured in filter sterilized Tryptic Soy broth. Both media were purchased from Becton, Dickinson and Company. Inocula for broth cultures were prepared by transferring a loopful of cells from a fresh agar plate into 200 μl of sterile saline, mixing the cells by vortex agitation, diluting the cells 1:1,000 in fresh broth, and then passing the inocula through a 5-μm pore-size syringe filter to remove large clumps of cells. Inocula ranged from 10^6^-10^7^ CFU/ml. Dual-species biofilms were inoculated at a *C. acnes*:*S. epidermidis* ratio of 2-8:1. Filtered inocula were aliquotied into sterile 13 × 100 mm glass tubes (1 ml/tube) and incubated at 37°C in air or in an anaerobic chamber. Anaerobic conditions were generated using a BD GasPak EZ Anaerobe sachet system. Tubes were incubated statically for 24 or 72 h, as indicated.

**TABLE 1.** Bacterial strains.

| Strain | Source | Characteristics | Reference |
| --- | --- | --- | --- |
| *Cutibacterium acnes* HL086PA1 | Acne patient | Ribotype RT8, Erm<sup>r</sup> | Fitz-Gibbon *et al.*, 2013* |
| *Staphylococcus epidermidis* 5 | Implant infection | PNAG-producing strain, Erm<sup>s</sup> | Chaignon *et al.*, 2007 |
| *Staphylococcus aureus* JE2 | Necrotizing fasciitis | MRSA, USA300 clone, Erm<sup>s</sup> | Miller *et al.*, 2005 |

\**C. acnes* HL086PA1 was obtained through BEI Resources as part of the NIAID NIH Human Microbiome Project.

### Crystal violet binding assay

Biofilms were rinsed vigorously with tap water to remove loosely adherent cells and stained for 1 min with 1 ml of Gramtis crystal violet. Tubes were then rinsed with tap water to remove the unbound dye, air-dried, and photographed. To quantitate crystal violet binding, stained tubes were filled with 1 ml of 33% acetic acid, incubated at room temperature for 30 min, and mixed by vortex agita6on. A volume of 200 μl of the dissolved dye was transferred to the well of a 96-well microtiter plate and its absorbance was measured in a microplate reader set to 620 nm. Tubes containing sterile broth were incubated and processed along with the inoculated tubes to serve as controls.

### Quantitation of biofilm CFUs

Biofilms were rinsed three times with 3 ml of saline and the tubes were filled with 1 ml of saline. Biofilm bacteria were detached from the walls of the tubes by sonication for 30 sec using an IKA Labortechnik sonicator set to 50% power and 50% duty cycle. Control experiments showed that this sonication treatment did not affect the viability of *C. acnes, S. epidermidis*, or *S. aureus* cells (data not shown). Sonicates were serially diluted in saline. To enumerate staphylococci, dilutions were plated on Tryp-c Soy agar and incubated in air. To enumerate *C. acnes*, dilutions were plated on Tryptic Soy agar supplemented with 20 μg/ml erythromycin and incubated anaerobically.

### Treatment of biofilms with enzymes

Biofilms were rinsed vigorously with water and then treated with 1 ml of 100 μg/ml dispersin B (Kane Biotech) or 100 μg/ml bovine DNase I (Sigma-Aldrich) in saline. Atier 15 min at 37°C, tubes were rinsed vigorously with water and stained with crystal violet as described above.

### Statistics and reproducibility of results

All experiments were performed in triplicate tubes. All experiments were performed on at least three occasions with similar results. The significance of differences between means was calculated using a Studenttis *t*-test. A *P-*value < 0.01 was considered significant.

## RESULTS

### *S. epidermidis* enhances *C. acnes* biofilm growth under anaerobic conditions and enables *C. acnes* to form biofilms under aerobic conditions

*S. epi-der-midis* and *C. acnes* were inoculated individually or jointly into broth in glass test tubes. The tubes were incubated at 37°C under anaerobic or aerobic conditions. Atter 72 h, the tubes were rinsed to remove planktonic and loosely adherent cells, and the biofilm cells were detached by sonication and enumerated by dilution plating (Fig. 1). When cultured individually, *S. epidermidis* formed biofilms under both anaerobic and aerobic conditions, whereas *C. acnes* formed biofilms under anaerobic conditions but not under aerobic conditions (Fig. 1, green and blue lines, respectively). *S. epidermidis* formed significantly more biofilm under aerobic conditions than under anaerobic conditions (Fig. 1, le2 panel, green and blue lines, respectively). When *S. epidermidis* and *C. acnes* were cultured together, the presence of *C. acnes* had no effect on the growth of *S. epidermidis* biofilms under either anaerobic or aerobic conditions (Fig. 1, le4 panel, violet and red lines, respectively). In contrast, the presence of *S. epidermidis* significantly enhanced the growth of *C. acnes* biofilms under anaerobic conditions and enabled *C. acnes* to form biofilms under aerobic conditions (Fig. 1, right panel, violet and red lines, respectively).

**Figure 1.**
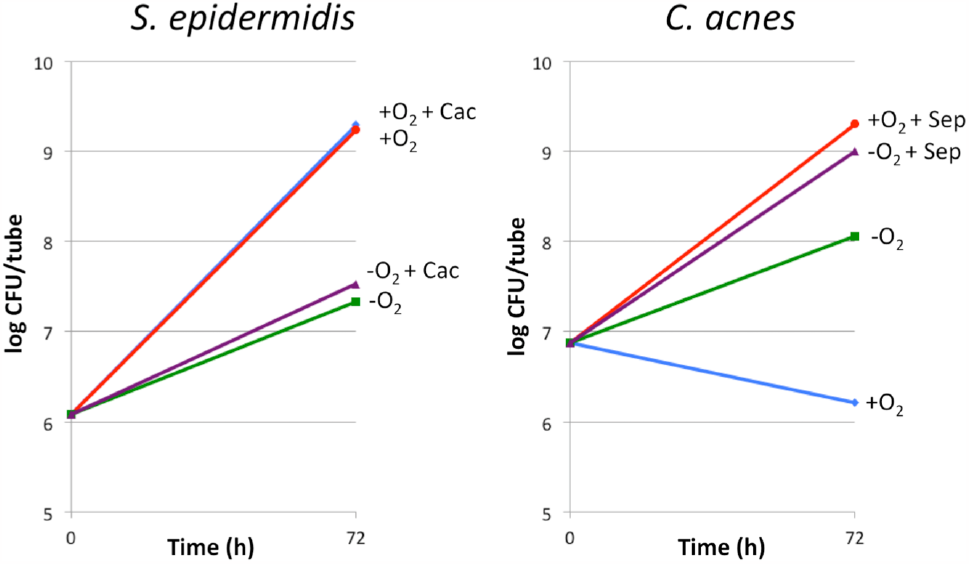
Growth of *S. epidermidis* (leti panel) and *C. acnes* (right panel) in monospecies and dual-species biofilms cultured under aerobic and anaerobic condi2ons. Blue line, mono-species biofilm aerobic (tiO_2_); green line, mono-species biofilm anaerobic (-O_2_); red line, dual-species biofilm aerobic; violet line, dual-species biofilm anaerobic. Cac, *C. acnes*; Sep, *S. epidermidis*. Values show means for triplicate tubes for each condition.

### Exopolysaccharide mediates *S. epidermidis/C. acnes* dual-species biofilm formation under aerobic conditions

Previous studies showed that the exopolysaccharide poly-*N*-acetyl-glucosamine (PNAG) and extracellular double-stranded DNA (eDNA) function as adhesive components in *S. epidermidis* and *C. acnes* biofilms (Kaplan *et al*., 2011; Kaplan, 2023). To determine whether PNAG or eDNA contribute to biofilm cohesion in aerobic *S. epidermidis*/*C. acnes* dual-species biofilms, biofilms were treated with the PNAG-degrading enzyme dispersin B (Kaplan *et al*., 2004), or with DNase I (Fig. 2). Treatment of dual-species biofilms with dispersin B, but not DNase I, resulted in a significant decrease in crystal violet stainable biomass (Figs. 2 & 3), suggesting that PNAG, but not eDNA, contributes to dual-species biofilm cohesion under the conditions tested.

**Figure 2.**
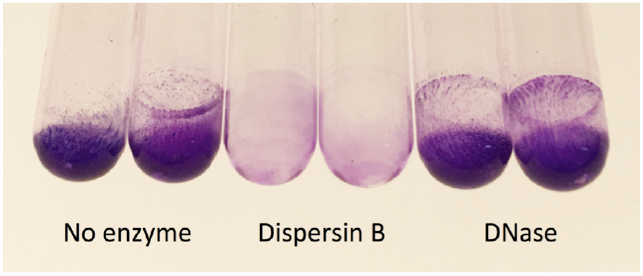
Detachment of aerobic *S. epidermidis/C. acnes* dual-species biofilms by dispersin B and DNase. Tubes were incubated for 24 h, rinsed with water, treated with the inducated enzyme for 15 min, re-reinsed, and stained with crystal violet. Duplicate tubes for each condition are shown.

**Figure 3.**
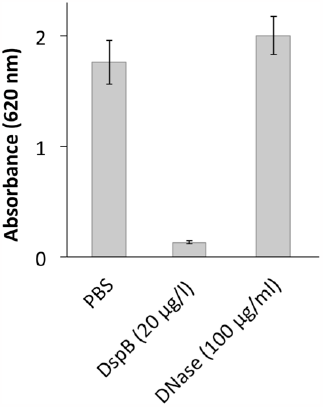
Quantitation of detachment of aerobic *S. epidermidis/C. acnes* dual-species biofilms by dispersin B and DNase. Values show mean and range for triplicate tubes for each condition.

### *S. aureus* enables *C. acnes* to form biofilms under aerobic conditions

*C. acnes* also formed biofilms under aerobic conditions when co-cultured with *S. aureus* JE2, a biofilm-forming MRSA strain (Fig. 4). This finding suggests that the biofilm enabling phenotype exhibited by *S. epidermidis* may be a general phenomenon and not be due to a specific interaction between *S. epidermidis* and *C. acnes*.

**Figure 4.**
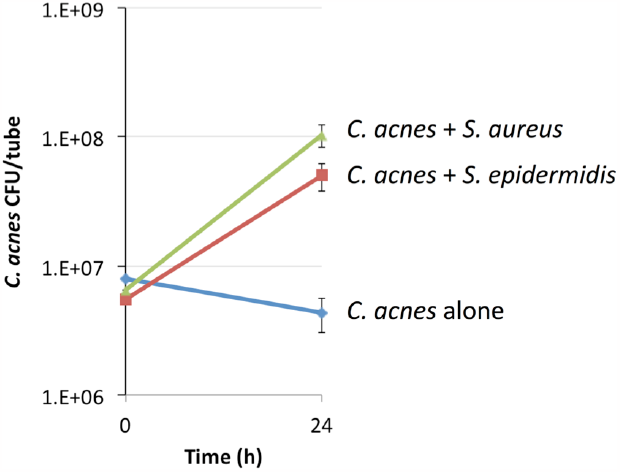
Aerobic growth of *C. acnes* in a mono-species biofilm (blue line), in a dual-species biofilm with *S. epidermidis* (red line), and in a dual-species biofilm with *S. aureus* (green line). Values show mean and range for triplicate tubes for each condition.

## DISCUSSION

Numerous studies have shown that under aerobic conditions, the growth of aerobic or facultatively anaerobic bacteria or fungi reduces the level of oxygen and thereby facilitates the growth of strictly anaerobic bacteria both in the planktonic and biofilm modes of growth. For example, a mixed culture of 10 bacterial species that included both aerobes and anaerobes was shown to protect obligate anaerobes from the toxic effects of oxygen when cultured in an aerated chemostat (Bradshaw *et al*., 1996). Similarly, the aerotolerant anaerobe *Fusobacterium nucleatum* was shown to enhance growth the strict anaerobe *Porphyromonas gingivalis* when cocultured in an aerated chemostat in which *P. gingivalis*, as a monoculture, was unable to survive (Diaz *et al*., 2002). Co-culture of *Enterococcus faecalis* with *P. gingivalis* supported aerobic growth of *P. gingivalis* in polystyrene micro.ter plates (Tan *et al*., 2022). Also, under aerobic conditions, the yeast-like fungus *Candida albicans* was shown to induce survival and proliferation of the anaerobic bacterial genera *Veillonella, Leptotrichia, Prevotella*, and *Fusobacterium* when added to human saliva and grown on glass coverslips (Janus *et al*., 2017); to increase the growth of the anaerobes *Clostridium perfringens* and *Bacteroides fragilis* when co-cultured in polystyrene microtiter plates (Fox *et al*., 2014); to increase the viability of *P. gingivalis* when co-cultured in polystyrene microtiter plates (Bartnickaet al., 2019); and to enhance the growth of *C. acnes* when co-cultured on polystyrene coupons (Bernard *et al*. 2019).

Ours is the first study to demonstrate that *S. epidermidis* enables *C. acnes* to form biofilms under oxic conditions. Our results also show that *S. epidermidis* significantly enhances biofilm growth of *C. acnes* under anaerobic conditions. In fact, when co-cultured with *S. epidermidis, C. acnes* biofilm formation was approximately equivalent under aerobic and anaerobic conditions. These phenotypes may result from decreased oxygen tension due to oxygen utilization by *S. epidermidis* during aerobic respiration, the formation of a protective biofilm niche by *S. epidermidis* that shields *C. acnes* from the inhibitory effects of oxygen, or the production of a factor by *S. epidermidis* that increases *C. acnes* aerotolerance or growth rate. The fact that *S. aureus* also enabled *C. acnes* to form biofilms under aerobic conditions suggests that the biofilm enabling phenotype may be due to non-specific interactions. Interestingly, *C. acnes* strain HL086PA1 did not form robust mono-species biofilms in glass tubes under anaerobic conditions when the biofilms were visualized using a crystal violet binding assay (Kaplan, 2023). These results suggest that *S. epidermidis* may enable biofilm formation even by non-bio-film-forming *C. acnes* strains. Another important finding was that PNAG and not eDNA functions as the major adhesion in mature *S. epidermidis*/*C. acnes* dual-species biofilms. More studies are needed to understand the interaction between *S. epidermidis* and *C. acnes* in dual-species and multi-species biofilms. The biofilm enabling phenotype may have clinical relevance to the interaction between *S. epidermidis* and *C. acnes* on the skin.

## ACKNOWLEDGEMENTS

The author thanks BEI Resources (Manassas, Virginia, USA) for providing *C. acnes* strain HL086PA1; Kane Biotech Inc. (Winnipeg, Manitoba, Canada) for providing dispersin B; and Katie DeCicco-Skinner (American University) for continued support.

## FUNDING

This project was funded by Kane Biotech Inc.

